# Cheetah: a computational toolkit for cybergenetic control

**DOI:** 10.1101/2020.06.25.171751

**Authors:** Elisa Pedone, Irene de Cesare, Criseida G. Zamora-Chimal, David Haener, Lorena Postiglione, Antonella La Regina, Barbara Shannon, Nigel J. Savery, Claire S. Grierson, Mario di Bernardo, Thomas E. Gorochowski, Lucia Marucci

**Affiliations:** Department of Engineering Mathematics, University of Bristol, Ada Lovelace Building, University Walk, BS8 1TW, Bristol, UK; School of Cellular and Molecular Medicine, University of Bristol, Biomedical Sciences Building, University Walk, BS8 1TD, Bristol, UK; BrisSynBio, Life Sciences Building, Tyndall Avenue, BS8 1TQ, Bristol, UK; School of Biochemistry, University of Bristol, Biomedical Sciences Building, University Walk, BS8 1TD, Bristol, UK; School of Biological Sciences, University of Bristol, Tyndall Avenue, BS8 1TQ, Bristol, UK; Department of EE and ICT, University of Naples Federico II, Via Claudio 21, 80125, Naples, Italy

**Author notes:** Correspondence should be addressed to T.E.G. and L.M. These authors contributed equally to this work and should be considered co-first authors (Elisa Pedone, Irene de Cesare, Criseida G. Zamora-Chimal, David Haener). These authors contributed equally to this work and should be considered co-senior authors (Thomas E. Gorochowski, Lucia Marucci).

**Keywords:** image analysis, microscopy, deep learning, U-Net, cybergenetics, synthetic biology

## Abstract

Advances in microscopy, microfluidics and optogenetics enable single-cell monitoring and environmental regulation and offer the means to control cellular phenotypes. The development of such systems is challenging and often results in bespoke setups that hinder reproducibility. To address this, we introduce Cheetah – a flexible computational toolkit that simplifies the integration of real-time microscopy analysis with algorithms for cellular control. Central to the platform is an image segmentation system based on the versatile U-Net convolutional neural network. This is supplemented with functionality to robustly count, characterise and control cells over time. We demonstrate Cheetah’s core capabilities by analysing long-term bacterial and mammalian cell growth and by dynamically controlling protein expression in mammalian cells. In all cases, Cheetah’s segmentation accuracy exceeds that of a commonly used thresholding-based method, allowing for more accurate control signals to be generated. Availability of this easy-to-use platform will make control engineering techniques more accessible and offer new ways to probe and manipulate living cells.

## Introduction

Modern automated microscopy techniques enable researchers to collect vast amounts of single-cell imaging data at high temporal resolutions. This has resulted in time-lapse microscopy becoming the go to method for studying cellular dynamics, enabling the quantification of processes such as stochastic fluctuations during gene expression ^1–3^, emerging oscillatory patterns in protein concentrations ^4^, lineage selection ^5,6^, and many more ^7^.

To make sense of microscopy images, segmentation is performed whereby an image is broken up into regions corresponding to specific features of interest (e.g. cells and the background). Image segmentation allows for the accurate quantification of cellular phenotypes encoded by visual cues (e.g. fluorescence) by ensuring only those pixels corresponding to a cell are considered. A range of segmentation algorithms have been proposed to automatically analyse images of various organisms and tissues ^3,8–11^. The most common of these are thresholding ^12^ and seeded watershed ^13^ methods, which are available in many scientific image processing toolkits. Commercial software packages also implement this type of functionality, enabling both automated image acquisition and analysis (e.g. NIS-Elements, Nikon). While these proprietary systems are user-friendly requiring no programming skills to be used, they are often difficult to tailor for specific needs and cannot be easily extended to new forms of analysis.

More recently, deep learning-based approaches to image segmentation have emerged ^7,14–17^. Compared to the more common thresholding-based approaches ^12^, deep learning methods tend to require more significant computational resources when running on traditional computer architectures, and often require the time-consuming manual step of generating large numbers of classified images for training. However, once trained deep learning methods are generally more robust to varying image quality and provide comparable ^18^ or superior segmentation accuracy ^17^ to thresholding-based methods.

The accuracy and robustness of a segmentation method are particularly important for online applications. For example, where an environment is dynamically controlled during an experiment in response to changes in cell state. Real-time image analysis and segmentation allow for the implementation of external feedback control ^19,20^. Typically, in such an experiment a combined microfluidic and microscopy platform is used to allow for images of single cells to be continually captured and analysed, with changes immediately processed. The state of the cells is generally signalled by the expression of a fluorescence protein that can be dynamically monitored and used as input to a control algorithm. The comparison of this cellular signal to the desired reference *in silico* allows a control signal to be generated by computer software that can be used to alter the cellular environment and perturb the cellular state in the required way (closing the loop). Generally, these experiments require the cells to be genetically engineered to transmit their state using fluorescence and respond to specific environmental stimuli in a prescribed way. This combination of computational, physical, and genetic aspects has resulted in this type of approach being termed “external cybergenetic control” and has been successfully applied for regulating gene expression and intracellular signalling in yeast ^21–26^, bacteria ^27^ and mammalian cells ^28–30^. Such external feedback control can also be implemented using optogenetics ^2,31^ and in combination with flow cytometry for online measurement of the control output ^32^. When compared to embedded cellular controllers (where both the controlled process and the controller are implemented within the cell using synthetic regulatory networks), external controllers benefit from requiring only minimal cellular modification, placing little burden on a cell; also, a single control platform can be used for the automatic regulation of different cellular processes across cellular species (e.g. gene expression ^21,22^, cell growth ^32^, cytosol-nuclear protein translocation ^33^).

In terms of software, while control algorithms such as proportional integral, model predictive control and zero average dynamics are versatile enough to be used in many contexts^34^, an online segmentation algorithm usually needs to be tailored given the cell type and the image acquisition settings. For example, if using a thresholding-based approach, various parameters in the segmentation code must be adjusted by trial-and-error before running a closed-loop control experiment. Furthermore, these settings must not significantly change during an experiment (e.g. due to a loss of focus), otherwise accuracy will be compromised. If the online measurements deviate from the real state of the cells, the overall control experiments will fail as inputs become calibrated to a miscalculated control error. Segmentation accuracy and robustness are therefore crucial for any form of control algorithm used.

In this work, we aim to address these difficulties by developing a computational toolkit called Cheetah to help simplify external cybergenetic control applications. We demonstrate its core functionality and flexibility by both post-processing time-lapse data for bacterial and mammalian cell growth in a microfluidic chip and external feedback control of gene expression in mammalian cells. Cheetah’s increased robustness is compared to the widely used Otsu thresholding-based method ^12,35^, here embedded in a multistep segmentation algorithm called ChipSeg ^33^, and we show how poor segmentation can lead to miscomputed control error and the possible failure of an experiment. Cheetah has a broad range of potential applications from post-experiment image analysis to robust real-time feedback control. Access to these capabilities in an easy-to-use package will help simplifying the integration of control engineering techniques into cell imaging platforms and offer new ways of robustly regulating the behaviour of living cells.

## Results and Discussion

### The Cheetah computational toolkit

Cheetah is a Python package designed to support closed-loop control in cybergenetic applications (i.e. systems that combine computational and genetic elements). It combines real-time image segmentation using the U-Net convolutional neural network (CNN) ^15,36^ with image analysis and cellular control algorithms. U-Net was chosen for segmentation because it has been proven reliable for a wide range of applications in systems and synthetic biology ^15,37,38^. Cheetah implements U-Net using Keras and to avoid overfitting, regularization can be customised to use either batch normalisation or a dropout rate (all examples in this work use batch normalisation).

Cheetah is composed of four modules (**Figure 1**). The first module supports the generation of training data for the U-Net model. Creation of training data can be laborious, therefore a ‘DataAugmentor’ class is provided to allow for a few labelled training images to be resampled and manipulated, generating a large, augmented set of training images. This works by sampling subregions of manually labelled images and then randomly applies image rotations, vertical and horizontal flips, scaling and shearing operations, and adjustments to the image histogram to simulate varying illumination levels. The use of these augmented training sets allows an accurate segmentation model to be trained using a far smaller number of manually labelled images ^15^.

**Figure 1:**
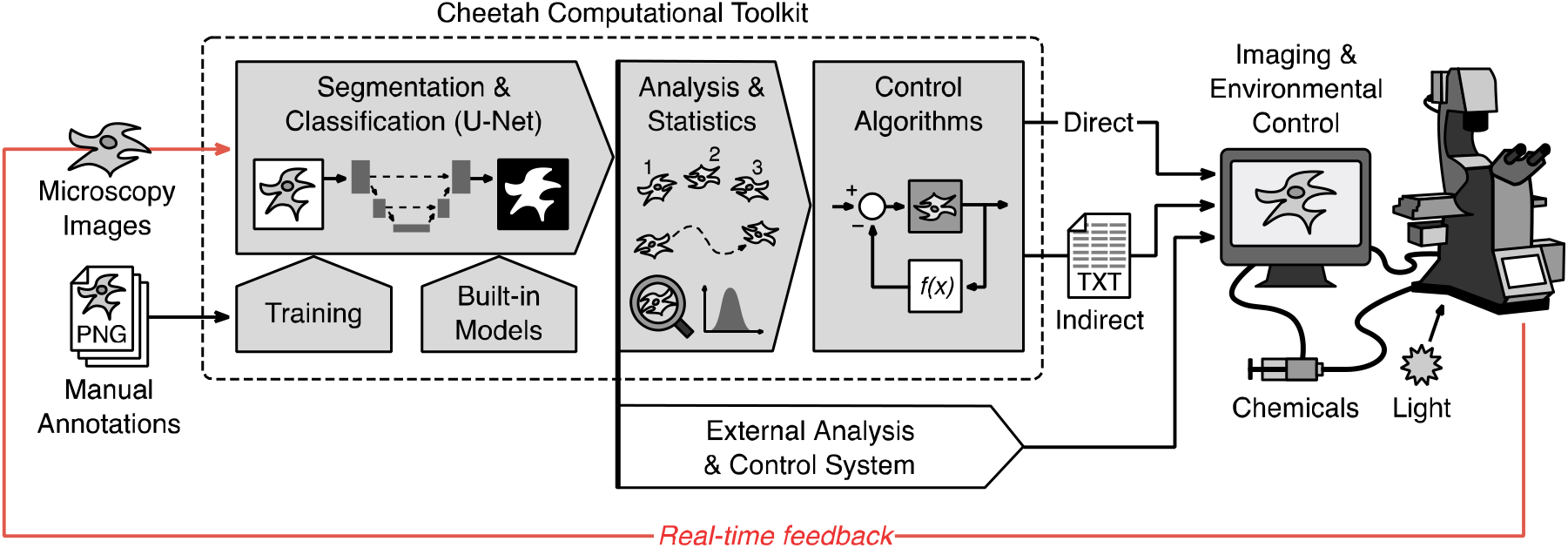
Overview of the Cheetah computational toolkit. Structure of Cheetah’s core modules and their interactions (grey filled arrows and boxes). The modular nature of the toolkit allows elements to be used separately, e.g., enabling the use of the built-in segmentation functionality with external analysis and control systems (white pointed box). Control algorithms can either directly interface with the imaging and environmental control system or output their data to text files for use by the external system (i.e. an indirect interface).

The second module is focused on the segmentation of images into various classes (e.g., class 1 = background, class 2 = cell). This functionality is defined within the ‘Segmenter’ class, which also includes functions to train the built-in U-Net model, to save and load the parameters for previously trained models, and to use a model for predicting the class of each pixel in a new image or image stack.

The third module takes segmented images as an input and can apply a range of common analyses. These include the extraction of pixel intensity histograms for a particular segmentation class (e.g. the intensity of all pixels within cells), the ability to classify and label separate cells, and to track cells across a time-series of images (provided movement is limited between frames).

Finally, the fourth module allows for the implementation of user-defined feedback control algorithms. These are implemented by extending the ‘ControlAlgorithm’ class, which includes placeholder functions for initialising the control setup and an execution loop that continually processes images and generates a control output that will be used to actuate the experimental setup. Built-in functions for Relay, Proportional–Integral (PI) and Proportional– Integral–Derivative (PID) control are provided as examples.

### Robust image segmentation and analysis of bacteria and mammalian cells

To demonstrate the core functionality of Cheetah, we made use of an integrated microfluidics and imaging platform that we have previously used for external feedback control of engineered bacterial and mammalian cells ^33^ (**Methods**). Previous time-lapse videos were collated and analysed using Cheetah and comparisons made to the same analyses performed using ChipSeg.

We began by post-processing an open-loop time-lapse experiment of *Escherichia coli* cells containing a genetic construct which uses an orthogonal σ/anti-σ pair to regulate expression of a green fluorescent protein (*gfp*) gene ^39^ (**Methods**). The experiment consisted of cells being grown in a microfluidic device designed for long-term bacterial culture ^40^ (**Figure 2A**) and images (including fluorescence) were acquired every 5 minutes over a 24-hour period (**Methods**). Before Cheetah could be used for analysis, it was necessary to train the system to be able to detect the bacteria in our experiment. This was done by manually annotating only 2 large images (512 × 512 pixels) containing 329 cells in total, with each pixel labelled as either ‘background’, ‘cell border’, or ‘cell interior’. The inclusion of the ‘cell border’ label was important to ensure separation of individual cells and accurate single cell analyses in densely packed regions (e.g. bacteria grown in a microfluidic chip) where ‘background’ pixels may not be visible between cells. These training images were augmented using Cheetah’s DataAugmentor class to create a final set of 60 smaller annotated images (256 × 256 pixels). Using this set of images allowed for a 99.2% segmentation accuracy to be reached after training, assessed using a random subset of images set aside solely for validation (**Methods**). Once trained, Cheetah segmentation masks were generated and used to calculate the number of cells and average GFP fluorescence per cell (**Figures 2B, 2C**). These results were compared to similar analyses using segmentation masks generated by ChipSeg that we ^33,41^ and others ^21,22^ have previously implemented in a similar experimental setup (**Supplementary Movie 1**; **Methods**).

**Figure 2:**
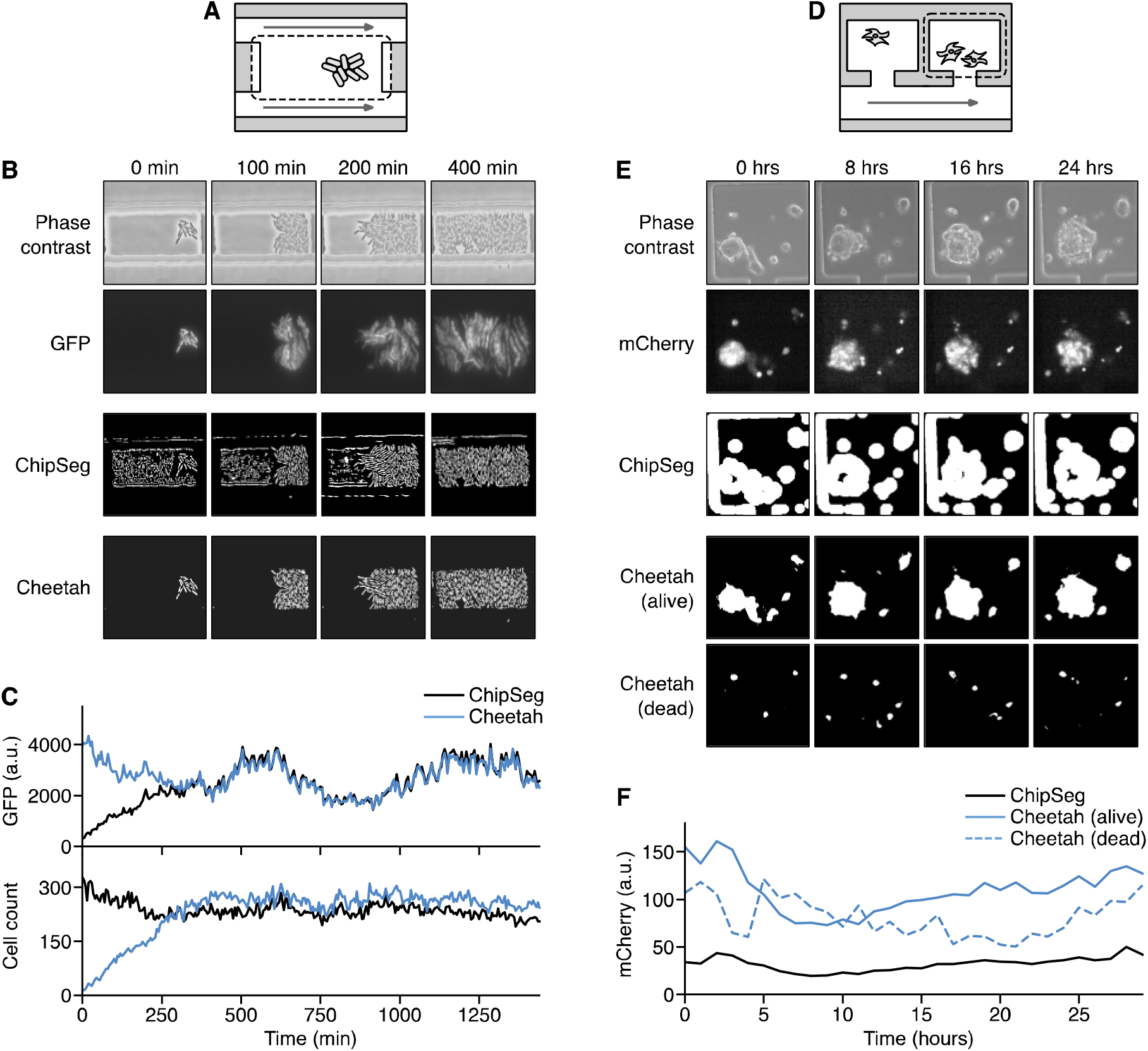
Monitoring and analysis of bacteria and mammalian cells in microfluidic chips. (**A**) Schematic of the microfluidic chamber used for bacterial growth and imaging. A typical imaging area is shown by the dashed box and flow of nutrients is shown by the grey arrows. (**B**) Time-lapse images of *Escherichia coli* cells growing in the microfluidic chamber for phase contrast and GFP fluorescence, as well as segmentation masks for cells generated using ChipSeg and Cheetah (white regions denote cells). (**C**) Average GFP fluorescence of the cell segmentation mask and cell count over time calculated using either ChipSeg or Cheetah segmentation masks. (**D**) Schematic of the microfluidic chamber used for mouse embryonic stem cells (mESCs) growth and imaging. A typical imaging area is shown by the dashed box and flow of nutrients is shown by the grey arrows. (**E**) Time-lapse images of mESCs growing in the microfluidic chamber for phase contrast and mCherry fluorescence, as well as segmentation masks for cells generated using ChipSeg and Cheetah (white regions denote cells). For Cheetah, separate masks are shown for living and dead cells. (**F**) Average mCherry fluorescence of the cell segmentation mask over time calculated using either ChipSeg or Cheetah.

There were several clear differences between the two segmentation methods. First, Cheetah gave more robust segmentation results, being able to accurately isolate the bacterial cells from their environment (**Figure 2B**). This differed from the ChipSeg segmentation, which struggled due to the edges of the microfluidic chamber and noise within the empty chamber caused by cell debris and fabrication imperfections that generated small high-contrast features. This resulted in the walls and empty regions of the chamber being recognised as cells and caused a large reduction in GFP fluorescence per cell at the start of the experiment when only a few cells were present (**Figure 2B**). As the experiment progressed, the impact of these misclassified regions was reduced as the majority of the image was covered in cells and so their impact was negligible. Furthermore, ChipSeg struggled to precisely distinguish individual cells, showing a visibly lower cell count once the chamber was filled with bacteria (**Figure 2C**). In contrast, Cheetah was not affected by any of these aspects and provided robust and reliable estimates of cell number and fluorescence per cell (**Figure 2C**) for the entire duration of the experiment. It should be noted that the significant difference of ~2600 arbitrary units (a.u.) in GFP fluorescence per cell at the beginning of the experiment between the methods would be a major problem for estimating a control signal, potentially causing large unwanted perturbations to the cells if used in an external feedback control system.

Bacterial cells generally have a simple and fairly consistent morphology across a population, which simplifies their classification. A more challenging problem is the analysis of mammalian cells whose shape can significantly vary over time. To assess Cheetah’s ability to handle these more complex cell types, we tested its ability to accurately isolate and characterise mouse embryonic stem cells (mESCs). Unlike in the bacterial example, mammalian cells can often die during an experiment, causing quantification of fluorescence to be influenced by these inactive cells. Ideally, dead cells should be excluded when calculating average fluorescence values, but often are not due to difficulties distinguishing each type with standard methods. Fortunately, this capability can be easily enabled in Cheetah due to the underlying U-Net segmentation model allowing for additional label types. Therefore, to analyse mammalian cells using Cheetah, we manually annotated 34 large images (1280 × 1056 pixels) containing 314 clones in total, with each pixel labelled as either ‘background’, ‘cell border’, ‘alive cell interior’, or ‘dead cell interior’ based on human knowledge regarding the generally smaller, disconnected and spherical shape of dead cells within a microfluidic chamber. Again, the DataAugmentor class was used to generate a final set of 536 smaller annotated images (512 × 512 pixels) which enabled Cheetah to reach a segmentation accuracy of 98.8% after training (**Methods**).

Next, we tested Cheetah using images from a 29-hour open-loop time-lapse experiment where engineered mESCs were grown in a microfluidic chamber that enabled long-term imaging (**Figure 2D**). mESCs were modified to carry an inducible genetic construct that expressed an mCherry fluorescent protein (**Methods**). As before, we compared the performance of cell segmentation and average mCherry fluorescence of Cheetah versus ChipSeg (**Supplementary Movie 2**). Consistent with the results for bacteria, ChipSeg incorrectly segmented the walls of the microfluidic chamber and struggled to precisely isolate cell bodies within the chamber (**Figure 2E**). When compared to the more accurate results generated by Cheetah, ChipSeg led to a slightly lower estimation of average mCherry fluorescence (**Figure 2F**). In contrast, Cheetah was able to classify alive and dead cells, which helped to improve its estimate of mCherry fluorescence for living cells; found to be marginally higher than for dead cells (**Figure 2F**).

### External feedback control of protein expression in mammalian cells

Having demonstrated the ability for Cheetah to robustly perform image analysis, we next attempted to validate its use for real-time external control of mammalian cells. Using the same engineered mESCs from the previous experiment, we employed an automated microscopy and fluidic control platform that allows for real-time live-cell imaging within microfluidic chips and the precise control of media and chemical inducers fed to the cells by the movement of motorised syringes (**Figure 3A**) ^28^. To allow for cells to be controlled by this system, mESCs carried a dual-input genetic construct where an mCherry fluorescent protein fused to a destabilising-domain (DD) was under the control of a ‘Tet-On’ promoter (**Figure 3B**, **Methods**) ^28^. This allowed the mCherry reporter to be switched ‘on’ by the combined presence of doxycycline (Doxy) and trimethoprim (TMP). By varying the concentration of these chemicals using the experimental platform in response to the deviation between the current mCherry fluorescence of the cells and the desired reference value, closed-loop real-time control of the cells could be achieved.

**Figure 3:**
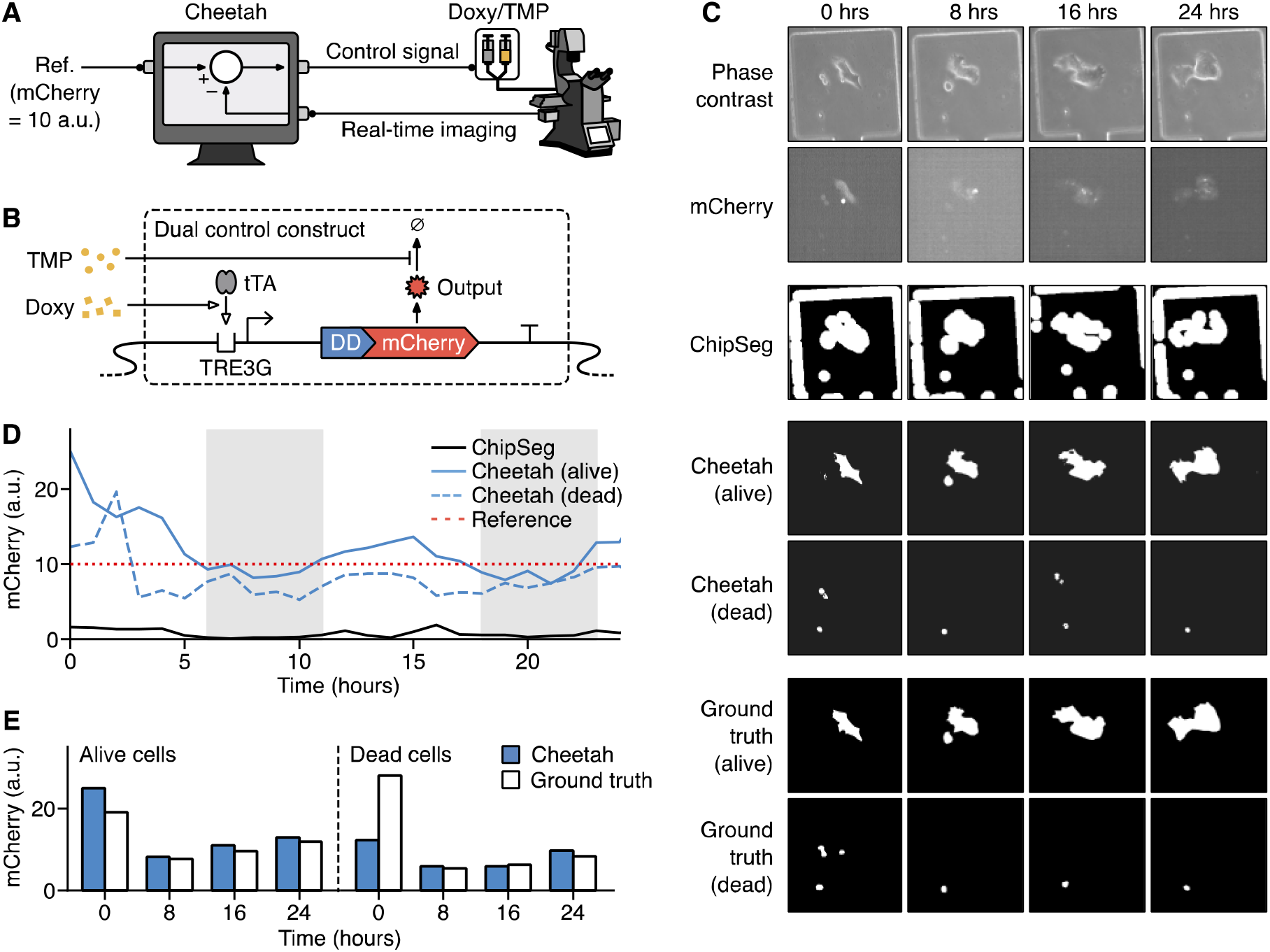
Closed-loop control of protein expression in mammalian cells. (**A**) Schematic of the microfluidic system used for external closed-loop control. A desired reference cellular mCherry fluorescence of 10 arbitrary units (a.u.) is shown. (**B**) Overview of genetic construct used to control mCherry expression ^28^. Small molecules (TMP and Doxy) work in tandem to boost the expression level of mCherry. Regulation is due to a tetracycline transcriptional activator (tTA) and a destabilising domain (DD) which forms part of the mCherry reporter protein. (**C**) Time-lapse images of mouse embryonic stem cells (mESCs) growing in the system for phase contrast and mCherry fluorescence, as well as segmentation masks for cells generated using ChipSeg, Cheetah and manually annotated to give a ground truth (white regions denote cells). For Cheetah and the ground truth, separate masks are shown for living and dead cells. (**D**) Average mCherry fluorescence of the cell segmentation mask over time calculated using either ChipSeg or Cheetah. Red dotted line denotes the external reference that the controller aims to maintain (10 a.u.) Grey shaded regions show when the control signal triggered release of TMP and Doxy. Control signals were generated by using average mCherry fluorescence calculated using segmentation masks of alive cells from Cheetah. (**E**) Comparison of average mCherry fluorescence at specific time points during the experiment for segmentation masks generated by Cheetah and manually annotated (ground truth).

To test the effectiveness of Cheetah for external *in silico* feedback control, mESCs carrying the dual-input genetic construct were exposed overnight to high concentrations of Doxy (1 μg/mL) and TMP (100 mM) to cause strong mCherry expression. These cells were then seeded into a microfluidic chip placed on our control platform (**Figure 3A**) and a Relay control algorithm ^28^ was used to allow for set-point regulation of mCherry expression over a period of 24 hours (**Methods**). In this case, we selected a desired reference average mCherry fluorescence of 10 arbitrary units (a.u.), which was half of the saturating mCherry fluorescence reached overnight and measured during the initialization phase of 120 minutes. For closed-loop feedback control, images were streamed to Cheetah every 60 minutes; each image was immediately segmented, and the mask generated for alive cells was used to estimate average mCherry fluorescence. This data was then fed to an external system to actuate the necessary control action (i.e. movement of the syringes and thus change in Dox and TMP concentration experienced by the cells) on the experimental platform.

Results from this experiment and related replica showed that the platform was able to accurately control average mCherry fluorescence in the cells for the duration of the experiment (**Figures 3C, 3D**; **Supplementary Figure 1**; **Supplementary Movie 3**). To evaluate the performance of the control experiment we measured the Integral Square Error (ISE) ^41^ for both controlled and uncontrolled chambers (i.e. those that received the same input in open loop). Across the three closed-loop experiments, we found that the ISE values were lower for living cells in the controlled chambers when compared to living cells in uncontrolled chambers that received the same input signals (**Supplementary Table 1**), confirming the effectiveness of our simple control strategy. As expected, within the control chambers, healthy living cells showed different ISEs as compared to those classified as dead or dying. For comparison, we analysed the same time-lapses data offline using ChipSeg (**Figure 3D, Supplementary Figure 1 C, D**; **Supplementary Movie 3**). Estimates of average mCherry fluorescence saw much lower levels due to misclassification of the chamber walls. Such incorrect estimation of fluorescence would have resulted in the mistaken triggering of the control input throughout the experiment.

To ensure that uncontrolled cells did not display similar behaviour as those exposed to varying inputs, experiments were performed in the presence and absence of the inducers Doxy and TMP (i.e. no chamber was controlled). As expected, these open loop experiments showed an increase or decay in mCherry fluorescence, respectively (**Supplementary Figures 2–4**).

We manually annotated 4 frames of the time-lapse data at 0, 8, 16, and 24 hours and compared the average mCherry fluorescence calculated using these masks and those automatically generated by Cheetah. This was done with no knowledge of the Cheetah segmentations to avoid bias. Close agreement was found for the alive and dead cells for most time points, with the only major deviation being for dead cells at 0 hours. Dead cells are often difficult to distinguish from living cells, so some differences, especially during seeding where cells are becoming accustomed to their new environment, would be expected (**Figure 3E**).

We also carried out a number of live/dead staining experiments using the fluorescent DNA stain 4’, 6-diamidino-2-phenylindole (DAPI) that accumulates in dead cells when present at high concentrations (**Methods**). These experiments also showed that Cheetah was able to accurately classify cells (**Supplementary Figure 5**) and suggest that the minor fluctuations we see in mCherry fluorescence for dead cells (**Figure 3D**) are due to small differences in the segmentation masks between timepoints and not due to further gene expression.

### Conclusions

As our ability to create cybergenetic systems that combine computational, physical, and biological elements advances, the need for supporting software to coordinate and control these systems will grow. Cheetah is an attempt to simplify this process by providing an easy-to-use computational toolkit that, while containing core functionality to speed up most projects, is also highly adaptable to new needs. Its major contribution is in providing a coherent computational framework that combines both robust U-Net image segmentation with analysis functions tailored for the rapid development of cybergenetic control setups. Here, we have demonstrated Cheetah’s abilities to rapidly segment and classify two morphologically different cell types in two different microfluidic settings. We show that Cheetah can rapidly compute highly accurate image segmentation (99.2% and 98.8% for *E. coli* and mESCs, respectively) even when trained using only a small number of manually annotated images (2 and 34 images for *E. coli* and mESCs, respectively). This exceeded the performance of ChipSeg ^12,35^ which achieved 96.1% and 74% accuracy for the same *E. coli* and mESCs data, respectively, and is comparable to other deep learning approaches (e.g. Delta which reaches 99.86% accuracy for bacteria when trained on a much larger set of images ^37^). Furthermore, we demonstrate how these capabilities allow for accurate control signals to be generated for external feedback control applications. In particular, the ability for Cheetah to not only segment, but also classify cells as potentially ‘dead’ or ‘alive’ enables it to filter out non-viable cells and leads to improved accuracy, as compared to non-classifier segmentation methods. In addition to segmentation and control algorithms, Cheetah also includes a wide range of built-in analysis for labelling cells, tracking their position across frames, and using this information to enable analysis of single-cell properties like fluorescence (see **Supplementary Movie 4** for an example). Being able to automate the creation of analysis dashboards, where multiple analyses are preformed and presented simultaneously, will also help speed up the discovery of subtle behaviours in populations of cells and offer the means to reanalyse existing time lapse microscopy data in more depth.

The ChipSeg algorithm that we used for comparison often performed poorly due to elements of the microfluidic chip (e.g. walls and imperfections in the fabrication) causing high contrast features that were incorrectly classified as cells. It should be noted that improved segmentation performance can be achieved through pre-processing of images to crop out unwanted features like the chamber walls ^35^. Such pre-processing was not performed here because for control applications it is often necessary to image over long-time courses, which can cause a drift in the images produced as many chambers are imaged sequentially. Drift in the images can make defining a cropping area difficult. This is especially true for small chambers where even small movements are a problem and for more complex microfluidic chip geometries such approaches may be impossible to reliably implement. Therefore, to be useful for cybergenetic control applications, the segmentation approaches must be able to handle images where aspects of the experimental setup are visible.

A potential disadvantage of deep learning approaches is the need to retrain the system when changes are made to the types of input data that will be received. For example, due to changes in an experimental setup or a different type of cell being used. While Cheetah’s use of data augmentation reduces the amount of manually annotated images required for training, there is growing interest in the area of transfer learning where a fully trained artificial neural network can be quickly adapted for effective use in a new, but related task ^42^. Given that other U-Net based segmentation systems have demonstrated the ability for transfer learning in other fields ^43^, this would be an interesting feature for future implementation in Cheetah.

To improve the quality and reusability of experimental data there has been growing interest in the use of external calibrants to enable the conversion of fluorescence ^44^ and gene expression levels ^45^ into calibrated or absolute units. In the context of imaging data, such conversions make it possible to directly compare measurements made between different microscopes and even labs, which would normally be impossible due to the arbitrary light intensity units that the images are captured in. Calibrated microscopy images could be produced by using fluorescently labelled beads to generate standard curves for conversion ^46^ and inclusion of functionality to interpret and convert images into calibrated units would be another useful future direction for this work.

While the focus here has been on demonstrating the major functionality of Cheetah, we anticipate that it can be applied much more broadly for applications across the field of synthetic biology. For example, using it within custom-built platforms able to perform imaging and dynamic light patterning ^47^ to control single-cells and guide collective behaviours ^48^. Furthermore, the code provided in the toolkit can easily be refined, customised, and extended to allow for new features to be implemented (e.g. more sophisticated control strategies). As such, Cheetah is a public, open-source project hosted on GitHub and welcomes contributions from the wider community.

We expect the deep learning methods that are central to Cheetah’s capabilities will play an increasingly important role in synthetic biology. In the context of external feedback control, the combination of deep learning-based label-free cell classification ^49,50^, online training approaches, model-free control strategies (e.g. reinforcement learning-based feedback control ^51^), and the availability of tunable genetic parts ^28,39,52,53^ could be instrumental in unlocking the potential for control engineering techniques in biology. This will open up new avenues to create reliable and robust, self-adaptive synthetic biological systems ^54^, much like how control engineering has revolutionised other fields.

## Methods

### Cheetah training process

Training of the U-Net convolutional neural network within Cheetah was performed using a Dell Precision 5530 laptop (Intel Core i7-8850H CPU, 16 GB RAM, and 512 GB NVMe SSD) running Windows 10, connected to a Sonnet eGFX Breakaway Box 550 hosting an NVIDIA Titan Xp GPU with 12 GB GDDR5X RAM. For all organisms, the full set of annotated images were randomly split with 70% used for training and the remaining 30% used for validation. Manual annotations used in the training for each organism were generated by a single person to avoid any differences in the classification of cell borders and live/dead cell classifications.

### ChipSeg segmentation

The Otsu segmentation method is based on pixel intensity levels and relies on the definition of grey threshold values used to divide a grayscale image into its components creating a binary mask ^12^. The simplest version of this algorithm allows for the identification of two-pixel classes, background and foreground, by using a single threshold level that aims to minimise the intra-class variance. More sophisticated versions of the algorithm couple global thresholding, previously described, to local thresholding, which computes dividing grey-intensity levels on smaller patches of the same image in order to boost the algorithm accuracy. In this work, segmentation was computed using ChipSeg ^33^, an Otsu-based method refined through the addition of both custom and MATLAB Image Processing Toolbox filtering functions. This pre-processing phase usually involves cropping smaller area of the original picture to be segmented and some filtering to increase saturation and resolution. Then, MATLAB functions implementing the Otsu method are executed, followed by the addiction of morphological operator to reshape the segmentation result. For further details and access to the code see de Cesare et al.^35^. For bacteria, both global and local thresholding was performed. The algorithm distinguished the foreground (single bacterial cells) from the background in each image of the time-lapse experiment. The global thresholding calculated the global area where cells are located, and the local thresholding found the centres and edges to differentiate individual cells in a binary mask. The final mask contained the boundaries and interiors of every segmented cell. This mask was overlaid to the fluorescence image field to calculate the fluorescence as the sum of all pixels in the segmented area minus the background fluorescence value. The average fluorescence across the bacterial population was then calculated as the mean of the fluorescence exhibited by all the objects in the final mask. Mammalian cells fluorescence was computed as the average pixel intensity value of pre-masked fluorescent images to which an average background intensity was subtracted, to take into account possible oscillations of microscopy’s light intensity. Masked images were obtained using the global thresholding strategy. It should be noted that background fluorescence values were similarly subtracted when using Cheetah.

### Bacterial strains, media and cell culture

Experiments with bacteria used a previously generated *E. coli* strain ^39^. Luria−Bertani (LB) medium (113002065, MP Biomedicals) supplemented with 50 μg/mL kanamycin (K4000, Sigma-Aldrich), 100 μg/mL ampicillin (A9518, Sigma-Aldrich) and 25 μg/mL chloramphenicol (C0378, Sigma-Aldrich) was used for all bacterial cell culture and microfluidics experiments. For microfluidic experiments, a single colony was used to seed 5 mL of LB media with antibiotics and grown overnight (approximately 16 hours) at 37°C with shaking at 200 rpm. 300 μL of the overnight culture was used to seed 300 mL of fresh LB medium with antibiotics. This culture was grown to an optical density at 600 nm of 0.3. The culture was then centrifuged at 2200 × *g* for 15 min and resuspended in 1.5 mL of fresh LB medium supplemented with 0.075% Tween-20 (P1379, Sigma-Aldrich) and antibiotics before loading into the microfluidic device.

### Mammalian cell lines, media and culture

Experiments with mammalian cells used a previously generated mouse Embryonic Stem Cell (mESC) line ^28^. Briefly, mESCs were subjected to two rounds of infection and drug-selection to stably express the transactivator (EF1a-rtTA, Neomycin) and the doxycycline-inducible vector (pLVX_TRE3GDDmCherry, Puromycin; Addgene plasmid #108679). Selected cells were expanded and grown on gelatin-coated dishes in Dulbecco’s modified Eagle’s medium (DMEM D5796, Sigma) supplemented with 15% fetal bovine serum (F7524, Sigma), 1X nonessential amino acids (11140035, Thermo Fisher), 2 mM L-Glutamine (25030024, Thermo Fisher), 100 μM 2-mercaptoethanol (31350010, Thermo Fisher), 1 mM Sodium Pyruvate (11360039, Thermo Fisher), 1X Penicillin/Streptomycin (P4458, Sigma) and 1000 U/mL LIF (250-02, Peprotech).

### Microfluidic devices and loading

For *E. coli*, the microfluidic device used was developed by Mondragón-Palomino and colleagues at the University of California, San Diego ^40^. A replica of the silicon mould was donated to our group. Soft lithography was used to form the microfluidic device which contains 48 trapping chambers and 6 inlet/outlet ports. Before each experiment, a wetting protocol was used to remove any air bubbles and debris from inside the device. The device was then mounted onto the stage of an inverted widefield fluorescence microscope, enclosed inside an incubation chamber set to 37°C (Pecon) and connected to fluidic lines. A cell loading protocol, trapping individual cells in the chambers of the device was performed via the C port. Ports W1 and W2 were used as waste ports, the C port became a waste port once the experiment had begun. Ports B and I were connected to an actuation system for motorised control of syringes to deliver fresh media and inputs to the cells growing inside the device. The R port was used as a mixing port. The microscope (see below for details) was programmed to take phase contrast (PhC), green fluorescence and red fluorescence images of the cells growing inside three different trapping chambers every 5 minutes. Green fluorescence images were used for the detection of sfGFP and red fluorescence images were used for the detection of the sulforhodamine B dye (230162, Sigma-Aldrich), used to detect the correct flow of inputs.

For mESCs, microfluidic chip loading and imaging were performed as reported previously ^28^. The microfluidic device we used was designed in the laboratory of Prof Jeff Hasty at the University California in San Diego. It consists of 5 ports for cell loading and media input/output, 33 individual chambers for cell growth and imaging, and a channel for controlled flow perfusion ^55^. The chip was fulfilled with complete mESC media supplemented with 1 μg/mL Doxy (D9891, Sigma) and 100 nM TMP (T7883, Sigma) flowing from port 5 followed by port 1 before the cell loading. Cells from a sub-confluent petri dish (60 cm in diameter) were washed with sterile Phosphate Buffered Saline (PBS) (D8537, Sigma), trypsinised for 2-3 min at room temperature and centrifuged at 1000 rpm for 5 min. Pelleted cells were resuspended in 200 μL of complete mESC medium+Doxy/TMP and gently loaded from port 1 using a 2 mL syringe, while applying constant vacuum suction to ports 3 and 4. The vacuum enables cell trapping by facilitating air release from the chambers. The chip was kept for 24 hours in a tissue culture incubator (5% CO_2_, 37°C) under constant Doxy/TMP perfusion to induce mCherry expression before the time-lapse. The day after, the device was transferred on the widefield microscope and connected to the actuation system that consists of two motor-controlled syringes (http://biodynamics.ucsd.edu/dialawave/) connected to port 6 and 7. One syringe contains Doxy, TMP and 1 μM of Atto488 green fluorescent dye (41051-1MG-F, ThermoFisher), whereas the other only contains plain mESC media. Ports 1, 2 and 5 were connected to stating syringes to balance the flow of media from ports 6 and 7, ensuring constant perfusion and avoiding backflow. During the open-loop experiments (**Figures 2E, F** and **Supplementary Figures 2-4**), mESCs were exposed to plain or Doxy/TMP supplemented media for the entire duration of the time-lapse, whereas dynamic switching between plain and Doxy/TMP media was automatically controlled in the closed-loop experiments (**Figures 3C, D** and **Supplementary Figure 1**) to reach and maintain a desired reference red fluorescence level.

### Live-cell imaging

Time-lapse microscopy for both *E. coli* and mESCs were performed using a Leica DMi8 inverted microscope equipped with an environmental control chamber (PeCon) for long-term temperature control and CO2 enrichment where necessary. The Adaptive Focus Control (AFC) ensures focus is maintained during the entire time-course experiment. Imaging of *E. coli* cells was performed using a 100X objective every 5 min using an AndoriXON 897 ultra back-illuminated EMCCD (512 × 512 pixel 16 μm pixels, 16-bit, 56 fps at full frame) in a temperature-controlled environment. Imaging of mESCs was performed using a 20X objective every 60 minutes in a temperature and CO_2_ controlled environment. The experimental set-up includes consecutive acquisition in three channels (phase contrast, green fluorescence and red fluorescence).

### Live/dead cell staining

To assess whether cells were alive, we performed experiments using the fluorescent DNA strain 4’, 6-diamidino-2-phenylindole (DAPI) (1ug/mL; D9542, Sigma) to stain on-chip mESCs before and after a microfluidic/microscopy-based time-lapse. When present at high concentrations, DAPI collects in dead cells. The accuracy of the Cheetah live/dead cell classifications were assessed by comparing the live/dead segmentation masks (generated purely from widefield images) to the fluorescence microscopy images of the stain (**Supplementary Figure 5**).

### Relay control algorithm

The Relay Control algorithm provides at each timepoint a control action that aims to minimise the error signal (*e*, defined as the difference between a reference signal and the process output). Formally, the controller generates the following control input

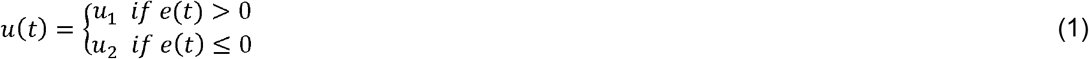

to decrease the error. In our experiments, the control input *u*_1_ corresponds to providing cells culture media supplemented with Doxy/TMP, while *u*_2_ corresponds to providing cells with plain media. The algorithm also implements a 5% hysteresis interval around the set-point to avoid chattering in the control signal.

### Integral Square Error (ISE)

To evaluate the performance of the controller, we used the integral square error (ISE), which is calculated as

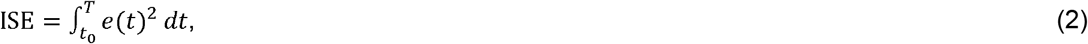

where *e*(*t*) is the difference between the reference mCherry fluorescence level and the measured mCherry fluorescence of the cells, and *t*_0_ and *T* are the start and end time of the experiment, respectively.

### General computational analysis and tools

Computational analysis was performed by custom scripts run using Python 3.6.8 and the following packages: tensorflow 1.14.0, keras 2.2.4, scikit-learn 0.21.2, scikit-image 0.15.0, numpy 1.16.4 and matplotlib 3.1.1. Genetic designs are visualised using DNAplotlib 1.0 ^56,57^ and Synthetic Biology Open Language Visual (SBOL Visual) symbols ^58^. Figures were composed using Omnigraffle 7.16 and Affinity Designer 1.8.3.

### Data availability

The Cheetah Python package, analysis code and data presented in this work are available from the project GitHub repository at: https://www.github.com/BiocomputeLab/cheetah.

## Supporting Information Statement

Supplementary Figure 1: Closed-loop control experiment; Supplementary Figures 2-4: Open-loop experiments; Supplementary Figure 5: Validation of cell classification; Supplementary Table 1: Integral Square Error (ISE); Supplementary Movie 1: Open-loop experiment of bacteria cells; Supplementary Movie 2: Open-loop experiment of mouse embryonic stem cells; Supplementary Movie 3: Closed-loop control experiment of mouse embryonic stem cells; Supplementary Movie 4: Analysis of on-chip growing bacteria cells

## Acknowledgements

T.E.G. gratefully acknowledges the support of NVIDIA Corporation for the donation of a Titan Xp GPU used in this research. This work was supported by BrisSynBio, a BBSRC/EPSRC Synthetic Biology Research Centre grant BB/L01386X/1 (T.E.G., L.M.), a Royal Society University Research Fellowship grant UF160357 (T.E.G.), the EU Horizon 2020 research project COSY-BIO grant 766840 (L.M.), EPSRC grants EP/R041695/1 and EP/S01876X/1 (L.M.), and MRC grant MR/N021444/1 (L.M.)

## Author contributions

L.M. and T.E.G. conceived of the project. D.H. developed the initial Keras implementation of the U-Net convolutional neural network. T.E.G. extended the U-Net implementation and developed the integrated Python package and cell analysis functions. E.P., C.Z., L.P., A.L.R. and B.S. performed experiments and generated training data. I.d.C. performed data analysis and supported computational aspects of the experimental work. L.M, T.E.G., N.S., C.S.G. and M.d.B. supervised the work. L.M. and T.E.G. wrote the paper with input from the other authors.

---

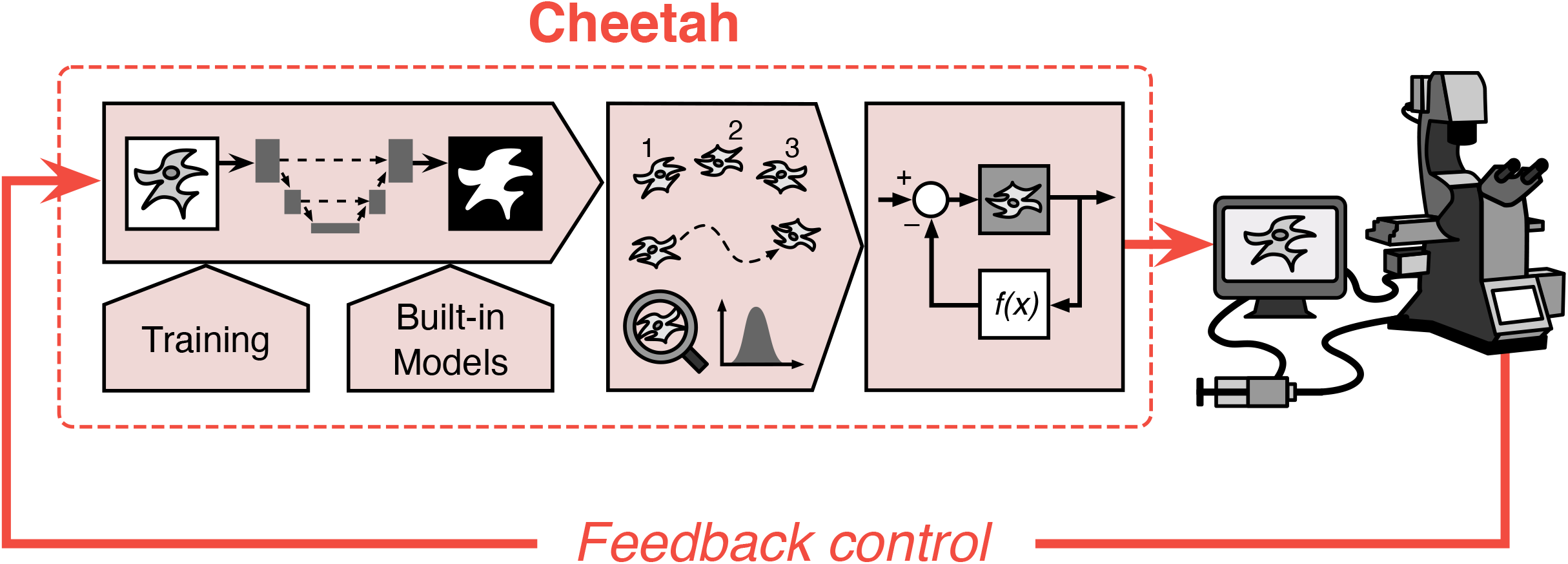

## References

(1) Marucci, L.; Pedone, E.; Di Vicino, U.; Sanuy-Escribano, B.; Isalan, M.; Cosma, M. P. β-Catenin Fluctuates in Mouse ESCs and Is Essential for Nanog-Mediated Reprogramming of Somatic Cells to Pluripotency. Cell Rep. 2014, 8 (6), 1686–1696. https://doi.org/10.1016/j.celrep.2014.08.011.

(2) Rullan, M.; Benzinger, D.; Schmidt, G. W.; Milias-Argeitis, A.; Khammash, M. An Optogenetic Platform for Real-Time, Single-Cell Interrogation of Stochastic Transcriptional Regulation. Mol. Cell 2018, 70 (4), 745–756.e6. https://doi.org/10.1016/j.molcel.2018.04.012.

(3) Young, J. W.; Locke, J. C. W.; Altinok, A.; Rosenfeld, N.; Bacarian, T.; Swain, P. S.; Mjolsness, E.; Elowitz, M. B. Measuring Single-Cell Gene Expression Dynamics in Bacteria Using Fluorescence Time-Lapse Microscopy. Nat. Protoc. 2012, 7 (1), 80–88. https://doi.org/10.1038/nprot.2011.432.

(4) Gritti, N.; Kienle, S.; Filina, O.; van Zon, J. S. Long-Term Time-Lapse Microscopy of C. Elegans Post-Embryonic Development. Nat. Commun. 2016, 7 (1), 12500. https://doi.org/10.1038/ncomms12500.

(5) Piltti, K. M.; Cummings, B. J.; Carta, K.; Manughian-Peter, A.; Worne, C. L.; Singh, K.; Ong, D.; Maksymyuk, Y.; Khine, M.; Anderson, A. J. Live-Cell Time-Lapse Imaging and Single-Cell Tracking of in Vitro Cultured Neural Stem Cells – Tools for Analyzing Dynamics of Cell Cycle, Migration, and Lineage Selection. Neural Stem Cells Health Dis. 2018, 133, 81–90. https://doi.org/10.1016/j.ymeth.2017.10.003.

(6) Brockway, N. L.; Cook, Z. T.; O’Gallagher, M. J.; Tobias, Z. J. C.; Gedi, M.; Carey, K. M.; Unni, V. K.; Pan, Y. A.; Metz, M. R.; Weissman, T. A. Multicolor Lineage Tracing Using in Vivo Time-Lapse Imaging Reveals Coordinated Death of Clonally Related Cells in the Developing Vertebrate Brain. Dev. Biol. 2019, 453 (2), 130–140. https://doi.org/10.1016/j.ydbio.2019.05.006.

(7) Bougen-Zhukov, N.; Loh, S. Y.; Lee, H. K.; Loo, L.-H. Large-Scale Image-Based Screening and Profiling of Cellular Phenotypes. Cytometry A 2017, 91 (2), 115–125. https://doi.org/10.1002/cyto.a.22909.

(8) E. Meijering. Cell Segmentation: 50 Years Down the Road. IEEE Signal Process. Mag. 2012, 29 (5), 140–145. https://doi.org/10.1109/MSP.2012.2204190.

(9) Stylianidou, S.; Brennan, C.; Nissen, S. B.; Kuwada, N. J.; Wiggins, P. A. SuperSegger: Robust Image Segmentation, Analysis and Lineage Tracking of Bacterial Cells. Mol. Microbiol. 2016, 102 (4), 690–700. https://doi.org/10.1111/mmi.13486.

(10) Kamentsky, L.; Jones, T. R.; Fraser, A.; Bray, M.-A.; Logan, D. J.; Madden, K. L.; Ljosa, V.; Rueden, C.; Eliceiri, K. W.; Carpenter, A. E. Improved Structure, Function and Compatibility for CellProfiler: Modular High-Throughput Image Analysis Software. Bioinformatics 2011, 27 (8), 1179–1180. https://doi.org/10.1093/bioinformatics/btr095.

(11) Eliceiri, K. W.; Berthold, M. R.; Goldberg, I. G.; Ibáñez, L.; Manjunath, B. S.; Martone, M. E.; Murphy, R. F.; Peng, H.; Plant, A. L.; Roysam, B. et al. Biological Imaging Software Tools. Nat. Methods 2012, 9 (7), 697–710. https://doi.org/10.1038/nmeth.2084.

(12) Otsu, N. A Threshold Selection Method from Gray-Level Histograns. IEEE Trans. Syst. Man Cybern. 1979, 9 (1), 62–66.

(13) Beucher, S.; Lantuejoul, C. Use of Watersheds in Contour Detection. In International Workshop on Image Processing: Real-Time Edge and Motion Detection/Estimation; Rennes, 1979.

(14) C. Sommer; C. Straehle; U. Köthe; F. A. Hamprecht. Ilastik: Interactive Learning and Segmentation Toolkit. In 2011 IEEE International Symposium on Biomedical Imaging: From Nano to Macro; 2011; pp 230–233. https://doi.org/10.1109/ISBI.2011.5872394.

(15) Falk, T.; Mai, D.; Bensch, R.; Çiçek, Ö.; Abdulkadir, A.; Marrakchi, Y.; Böhm, A.; Deubner, J.; Jäckel, Z.; Seiwald, K. et al. U-Net: Deep Learning for Cell Counting, Detection, and Morphometry. Nat. Methods 2019, 16 (1), 67–70. https://doi.org/10.1038/s41592-018-0261-2.

(16) Hilsenbeck, O.; Schwarzfischer, M.; Loeffler, D.; Dimopoulos, S.; Hastreiter, S.; Marr, C.; Theis, F. J.; Schroeder, T. FastER: A User-Friendly Tool for Ultrafast and Robust Cell Segmentation in Large-Scale Microscopy. Bioinformatics 2017, 33 (13), 2020–2028. https://doi.org/10.1093/bioinformatics/btx107.

(17) Caicedo, J. C.; Roth, J.; Goodman, A.; Becker, T.; Karhohs, K. W.; Broisin, M.; Molnar, C.; McQuin, C.; Singh, S.; Theis, F. J. et al. Evaluation of Deep Learning Strategies for Nucleus Segmentation in Fluorescence Images. Cytometry A 2019, 95 (9), 952–965. https://doi.org/10.1002/cyto.a.23863.

(18) Van Valen, D. A.; Kudo, T.; Lane, K. M.; Macklin, D. N.; Quach, N. T.; DeFelice, M. M.; Maayan, I.; Tanouchi, Y.; Ashley, E. A.; Covert, M. W. Deep Learning Automates the Quantitative Analysis of Individual Cells in Live-Cell Imaging Experiments. PLOS Comput. Biol. 2016, 12 (11), e1005177. https://doi.org/10.1371/journal.pcbi.1005177.

(19) M. Khammash; M. Di Bernardo; D. Di Bernardo. Cybergenetics: Theory and Methods for Genetic Control System. In 2019 IEEE 58th Conference on Decision and Control (CDC); 2019; pp 916–926. https://doi.org/10.1109/CDC40024.2019.9030209.

(20) Del Vecchio, D.; Dy, A. J.; Qian, Y. Control Theory Meets Synthetic Biology. J. R. Soc. Interface 2016, 13 (120), 20160380. https://doi.org/10.1098/rsif.2016.0380.

(21) Menolascina, F.; Fiore, G.; Orabona, E.; De Stefano, L.; Ferry, M.; Hasty, J.; di Bernardo, M.; di Bernardo, D. In-Vivo Real-Time Control of Protein Expression from Endogenous and Synthetic Gene Networks. PLOS Comput. Biol. 2014, 10 (5), e1003625. https://doi.org/10.1371/journal.pcbi.1003625.

(22) F. Menolascina; M. di Bernardo; D. di Bernardo. Design and Implementation of a Feedback Control Strategy for IRMA, a Novel Synthetic Gene Regulatory Network. In 49th IEEE Conference on Decision and Control (CDC); 2010; pp 2535–2540.

(23) Uhlendorf, J.; Miermont, A.; Delaveau, T.; Charvin, G.; Fages, F.; Bottani, S.; Batt, G.; Hersen, P. Long-Term Model Predictive Control of Gene Expression at the Population and Single-Cell Levels. Proc. Natl. Acad. Sci. 2012, 109 (35), 14271. https://doi.org/10.1073/pnas.1206810109.

(24) Perrino, G.; Wilson, C.; Santorelli, M.; di Bernardo, D. Quantitative Characterization of α-Synuclein Aggregation in Living Cells through Automated Microfluidics Feedback Control. Cell Rep. 2019, 27 (3), 916–927.e5. https://doi.org/10.1016/j.celrep.2019.03.081.

(25) Milias-Argeitis, A.; Summers, S.; Stewart-Ornstein, J.; Zuleta, I.; Pincus, D.; El-Samad, H.; Khammash, M.; Lygeros, J. In Silico Feedback for in Vivo Regulation of a Gene Expression Circuit. Nat. Biotechnol. 2011, 29 (12), 1114–1116. https://doi.org/10.1038/nbt.2018.

(26) Toettcher, J. E.; Gong, D.; Lim, W. A.; Weiner, O. D. Light-Based Feedback for Controlling Intracellular Signaling Dynamics. Nat. Methods 2011, 8 (10), 837–839. https://doi.org/10.1038/nmeth.1700.

(27) Lugagne, J.-B.; Sosa Carrillo, S.; Kirch, M.; Köhler, A.; Batt, G.; Hersen, P. Balancing a Genetic Toggle Switch by Real-Time Feedback Control and Periodic Forcing. Nat. Commun. 2017, 8 (1), 1671. https://doi.org/10.1038/s41467-017-01498-0.

(28) Pedone, E.; Postiglione, L.; Aulicino, F.; Rocca, D. L.; Montes-Olivas, S.; Khazim, M.; di Bernardo, D.; Pia Cosma, M.; Marucci, L. A Tunable Dual-Input System for on-Demand Dynamic Gene Expression Regulation. Nat. Commun. 2019, 10 (1), 4481. https://doi.org/10.1038/s41467-019-12329-9.

(29) Khazim, M.; Postiglione, L.; Pedone, E.; Rocca, D. L.; Zahra, C.; Marucci, L. Towards Automated Control of Embryonic Stem Cell Pluripotency. 8th Conf. Found. Syst. Biol. Eng. FOSBE 2019 2019, 52 (26), 82–87. https://doi.org/10.1016/j.ifacol.2019.12.240.

(30) Khazim, M.; Pedone, E.; Postiglione, L.; di Bernardo, D.; Marucci, L. A Microfluidic/Microscopy-Based Platform for on-Chip Controlled Gene Expression in Mammalian Cells. In Synthetic Gene Circuits◻: Methods and Protocols; Menolascina, F., Ed.; Springer US: New York, NY, 2021; pp 205–219. https://doi.org/10.1007/978-1-0716-1032-9_10.

(31) A. M. R. Denniss; T. E. Gorochowski; S. Hauert. Augmented Reality for the Engineering of Collective Behaviours in Microsystems. In 2019 International Conference on Manipulation, Automation and Robotics at Small Scales (MARSS); 2019; pp 1–6. https://doi.org/10.1109/MARSS.2019.8860907.

(32) Milias-Argeitis, A.; Rullan, M.; Aoki, S. K.; Buchmann, P.; Khammash, M. Automated Optogenetic Feedback Control for Precise and Robust Regulation of Gene Expression and Cell Growth. Nat. Commun. 2016, 7, 12546.

(33) Postiglione, L.; Napolitano, S.; Pedone, E.; Rocca, D. L.; Aulicino, F.; Santorelli, M.; Tumaini, B.; Marucci, L.; di Bernardo, D. Regulation of Gene Expression and Signaling Pathway Activity in Mammalian Cells by Automated Microfluidics Feedback Control. ACS Synth. Biol. 2018, 7 (11), 2558–2565. https://doi.org/10.1021/acssynbio.8b00235.

(34) Fiore, G.; Perrino, G.; di Bernardo, M.; di Bernardo, D. In Vivo Real-Time Control of Gene Expression: A Comparative Analysis of Feedback Control Strategies in Yeast. ACS Synth. Biol. 2016, 5 (2), 154–162. https://doi.org/10.1021/acssynbio.5b00135.

(35) de Cesare, I.; Zamora-Chimal, C. G.; Postiglione, L.; Khazim, M. R.; Pedone, E.; Shannon, B.; Fiore, G.; Perrino, G.; Napolitano, S.; di Bernardo, D. et al. ChipSeg: An Automatic Tool to Segment Bacteria and Mammalian Cells Cultured in Microfluidic Devices. ACS Omega 2020. https://doi.org/10.1021/acsomega.0c03906.

(36) Ronneberger, O.; Fischer, P.; Brox, T. U-Net: Convolutional Networks for Biomedical Image Segmentation. In Medical Image Computing and Computer-Assisted Intervention– MICCAI 2015; Navab, N., Hornegger, J., Wells, W. M., Frangi, A. F., Eds.; Springer International Publishing: Cham, 2015; pp 234–241.

(37) Lugagne, J.-B.; Lin, H.; Dunlop, M. J. DeLTA: Automated Cell Segmentation, Tracking, and Lineage Reconstruction Using Deep Learning. PLOS Comput. Biol. 2020, 16 (4), e1007673. https://doi.org/10.1371/journal.pcbi.1007673.

(38) Weigert, M.; Schmidt, U.; Boothe, T.; Müller, A.; Dibrov, A.; Jain, A.; Wilhelm, B.; Schmidt, D.; Broaddus, C.; Culley, S. et al. Content-Aware Image Restoration: Pushing the Limits of Fluorescence Microscopy. Nat. Methods 2018, 15 (12), 1090–1097. https://doi.org/10.1038/s41592-018-0216-7.

(39) Annunziata, F.; Matyjaszkiewicz, A.; Fiore, G.; Grierson, C. S.; Marucci, L.; di Bernardo, M.; Savery, N. J. An Orthogonal Multi-Input Integration System to Control Gene Expression in Escherichia Coli. ACS Synth. Biol. 2017, 6 (10), 1816–1824. https://doi.org/10.1021/acssynbio.7b00109.

(40) Mondragón-Palomino, O.; Danino, T.; Selimkhanov, J.; Tsimring, L.; Hasty, J. Entrainment of a Population of Synthetic Genetic Oscillators. Science 2011, 333 (6047), 1315. https://doi.org/10.1126/science.1205369.

(41) Shannon, B.; Zamora-Chimal, C. G.; Postiglione, L.; Salzano, D.; Grierson, C. S.; Marucci, L.; Savery, N. J.; di Bernardo, M. In Vivo Feedback Control of an Antithetic Molecular-Titration Motif in Escherichia Coli Using Microfluidics. ACS Synth. Biol. 2020, 9 (10), 2617–2624. https://doi.org/10.1021/acssynbio.0c00105.

(42) Kensert, A.; Harrison, P. J.; Spjuth, O. Transfer Learning with Deep Convolutional Neural Networks for Classifying Cellular Morphological Changes. SLAS Discov. Adv. Sci. Drug Discov. 2019, 24 (4), 466–475. https://doi.org/10.1177/2472555218818756.

(43) Adiba, A.; Hajji, H.; Maatouk, M. Transfer Learning and U-Net for Buildings Segmentation. Proceedings of the New Challenges in Data Sciences: Acts of the Second Conference of the Moroccan Classification Society, 2019, Article 14.

(44) Beal, J.; Haddock-Angelli, T.; Baldwin, G.; Gershater, M.; Dwijayanti, A.; Storch, M.; de Mora, K.; Lizarazo, M.; Rettberg, R.; with the iGEM Interlab Study Contributors. Quantification of Bacterial Fluorescence Using Independent Calibrants. PLOS ONE 2018, 13 (6), e0199432. https://doi.org/10.1371/journal.pone.0199432.

(45) Gorochowski, T. E.; Chelysheva, I.; Eriksen, M.; Nair, P.; Pedersen, S.; Ignatova, Z. Absolute Quantification of Translational Regulation and Burden Using Combined Sequencing Approaches. Mol. Syst. Biol. 2019, 15 (5), e8719. https://doi.org/10.15252/msb.20188719.

(46) Castillo-Hair, S. M.; Sexton, J. T.; Landry, B. P.; Olson, E. J.; Igoshin, O. A.; Tabor, J. J. FlowCal: A User-Friendly, Open Source Software Tool for Automatically Converting Flow Cytometry Data from Arbitrary to Calibrated Units. ACS Synth. Biol. 2016, 5 (7), 774–780. https://doi.org/10.1021/acssynbio.5b00284.

(47) Rubio Denniss, A.; Gorochowski, T. E.; Hauert, S. An Open Platform for High-Resolution Light-Based Control of Microscopic Collectives. bioRxiv 2020, 2020.12.28.424547. https://doi.org/10.1101/2020.12.28.424547.

(48) Gorochowski, T. E.; Hauert, S.; Kreft, J.-U.; Marucci, L.; Stillman, N. R.; Tang, T.-Y. D.; Bandiera, L.; Bartoli, V.; Dixon, D. O. R.; Fedorec, A. J. H. et al. Toward Engineering Biosystems with Emergent Collective Functions. Front. Bioeng. Biotechnol. 2020, 8, 705. https://doi.org/10.3389/fbioe.2020.00705.

(49) Chen, C. L.; Mahjoubfar, A.; Tai, L.-C.; Blaby, I. K.; Huang, A.; Niazi, K. R.; Jalali, B. Deep Learning in Label-Free Cell Classification. Sci. Rep. 2016, 6 (1), 21471. https://doi.org/10.1038/srep21471.

(50) Matek, C.; Schwarz, S.; Spiekermann, K.; Marr, C. Human-Level Recognition of Blast Cells in Acute Myeloid Leukaemia with Convolutional Neural Networks. Nat. Mach. Intell. 2019, 1 (11), 538–544. https://doi.org/10.1038/s42256-019-0101-9.

(51) Treloar, N. J.; Fedorec, A. J. H.; Ingalls, B.; Barnes, C. P. Deep Reinforcement Learning for the Control of Microbial Co-Cultures in Bioreactors. PLOS Comput. Biol. 2020, 16 (4), e1007783. https://doi.org/10.1371/journal.pcbi.1007783.

(52) Bartoli, V.; Meaker, G. A.; di Bernardo, M.; Gorochowski, T. E. Tunable Genetic Devices through Simultaneous Control of Transcription and Translation. Nat. Commun. 2020, 11 (1), 2095. https://doi.org/10.1038/s41467-020-15653-7.

(53) Baumschlager, A.; Aoki, S. K.; Khammash, M. Dynamic Blue Light-Inducible T7 RNA Polymerases (Opto-T7RNAPs) for Precise Spatiotemporal Gene Expression Control. ACS Synth. Biol. 2017, 6 (11), 2157–2167. https://doi.org/10.1021/acssynbio.7b00169.

(54) Bartoli, V.; di Bernardo, M.; Gorochowski, T. E. Self-Adaptive Biosystems through Tunable Genetic Parts and Circuits. Curr. Opin. Syst. Biol. 2020, 24, 78–85. https://doi.org/10.1016/j.coisb.2020.10.006.

(55) Kolnik, M.; Tsimring, L. S.; Hasty, J. Vacuum-Assisted Cell Loading Enables Shear-Free Mammalian Microfluidic Culture. Lab. Chip 2012, 12 (22), 4732–4737. https://doi.org/10.1039/C2LC40569E.

(56) Der, B. S.; Glassey, E.; Bartley, B. A.; Enghuus, C.; Goodman, D. B.; Gordon, D. B.; Voigt, C. A.; Gorochowski, T. E. DNAplotlib: Programmable Visualization of Genetic Designs and Associated Data. ACS Synth. Biol. 2017, 6 (7), 1115–1119. https://doi.org/10.1021/acssynbio.6b00252.

(57) Bartoli, V.; Dixon, D. O. R.; Gorochowski, T. E. Automated Visualization of Genetic Designs Using DNAplotlib. In Synthetic Biology: Methods and Protocols; Braman, J. C., Ed.; Springer New York: New York, NY, 2018; pp 399–409. https://doi.org/10.1007/978-1-4939-7795-6_22.

(58) Beal, J.; Nguyen, T.; Gorochowski, T. E.; Goñi-Moreno, A.; Scott-Brown, J.; McLaughlin, J. A.; Madsen, C.; Aleritsch, B.; Bartley, B.; Bhakta, S. et al. Communicating Structure and Function in Synthetic Biology Diagrams. ACS Synth. Biol. 2019, 8 (8), 1818–1825. https://doi.org/10.1021/acssynbio.9b00139.

