## Supplementary Information for "Cheetah: a computational toolkit for cybergenetic control"

### Table of Contents

| **Supplementary Figures** | **Page** |
|---|---|
| Supplementary Figure 1: Closed-loop control of protein expression in mammalian cells | 2 |
| Supplementary Figure 2: Open-loop protein expression in mammalian cells (rep. 1) | 3 |
| Supplementary Figure 3: Open-loop protein expression in mammalian cells (rep. 2) | 4 |
| Supplementary Figure 4: Open-loop protein expression in mammalian cells (rep. 3) | 5 |
| Supplementary Figure 5: Experimental verification of alive/dead cell classifications | 6 |
| **Supplementary Tables** | |
| Supplementary Table 1: Integral square errors for time-lapse segmentations | 7 |
| **Supplementary Movie Captions** | 8 |

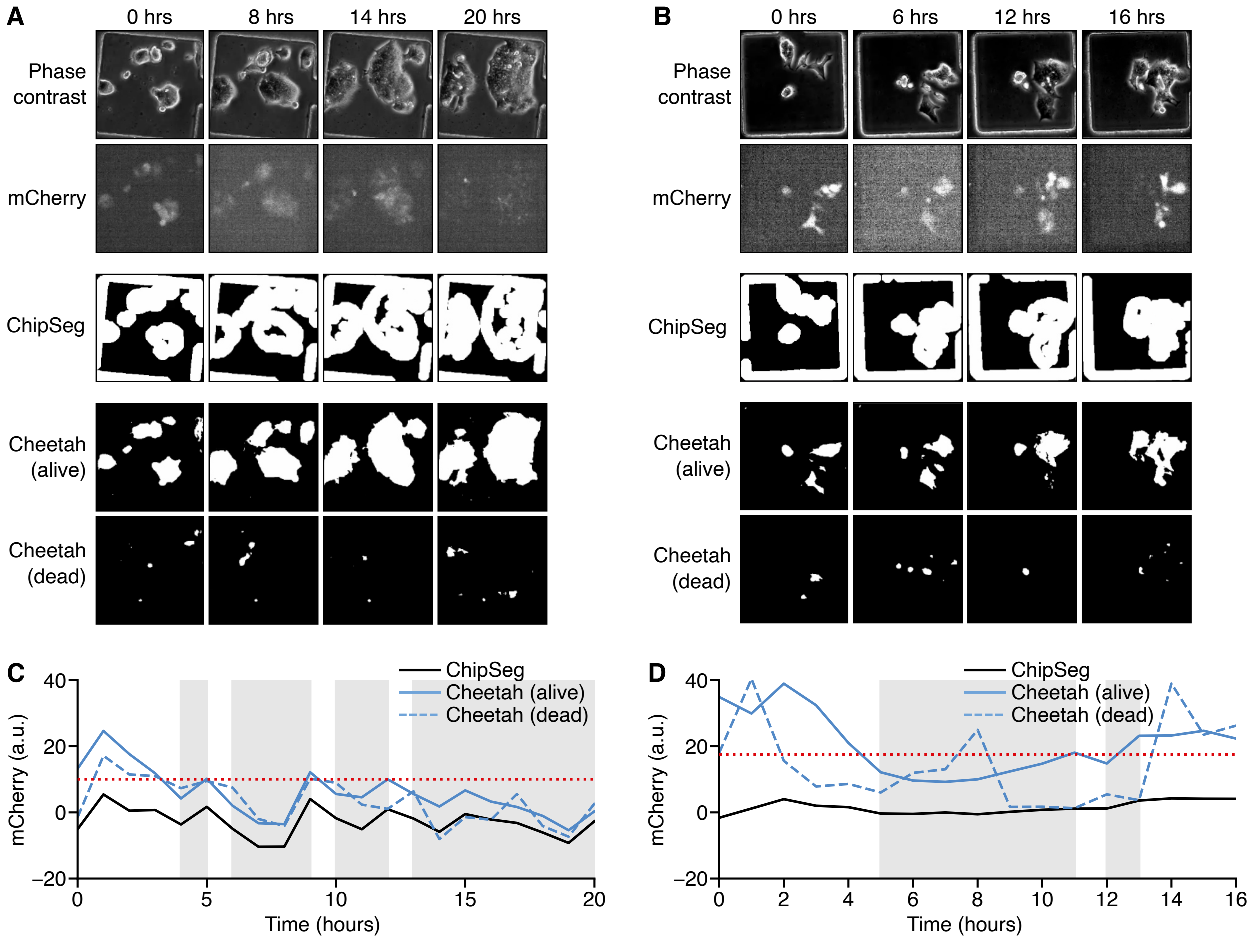


**Supplementary Figure 1: Closed-loop control of protein expression in mammalian cells.** Biological replicates of control experiment shown in Figure 3. (**A**, **B**) Time-lapse images of mouse embryonic stem cells (mESCs) growing in the system for phase contrast and mCherry fluorescence, as well as segmentation masks for cells generated using the Otsu method, and Cheetah. (**C**, **D**) Average mCherry fluorescence of the cell segmentation mask over time calculated using either ChipSeg or Cheetah. Red dotted line denotes the external reference that the controller aims to maintain (10 a.u. and 17 a.u. for panels C and D, respectively). Grey shaded regions show when the control signal triggered release of TMP and Doxy. Control signals were generated by using average mCherry fluorescence calculated from segmentation masks of alive cells from Cheetah.

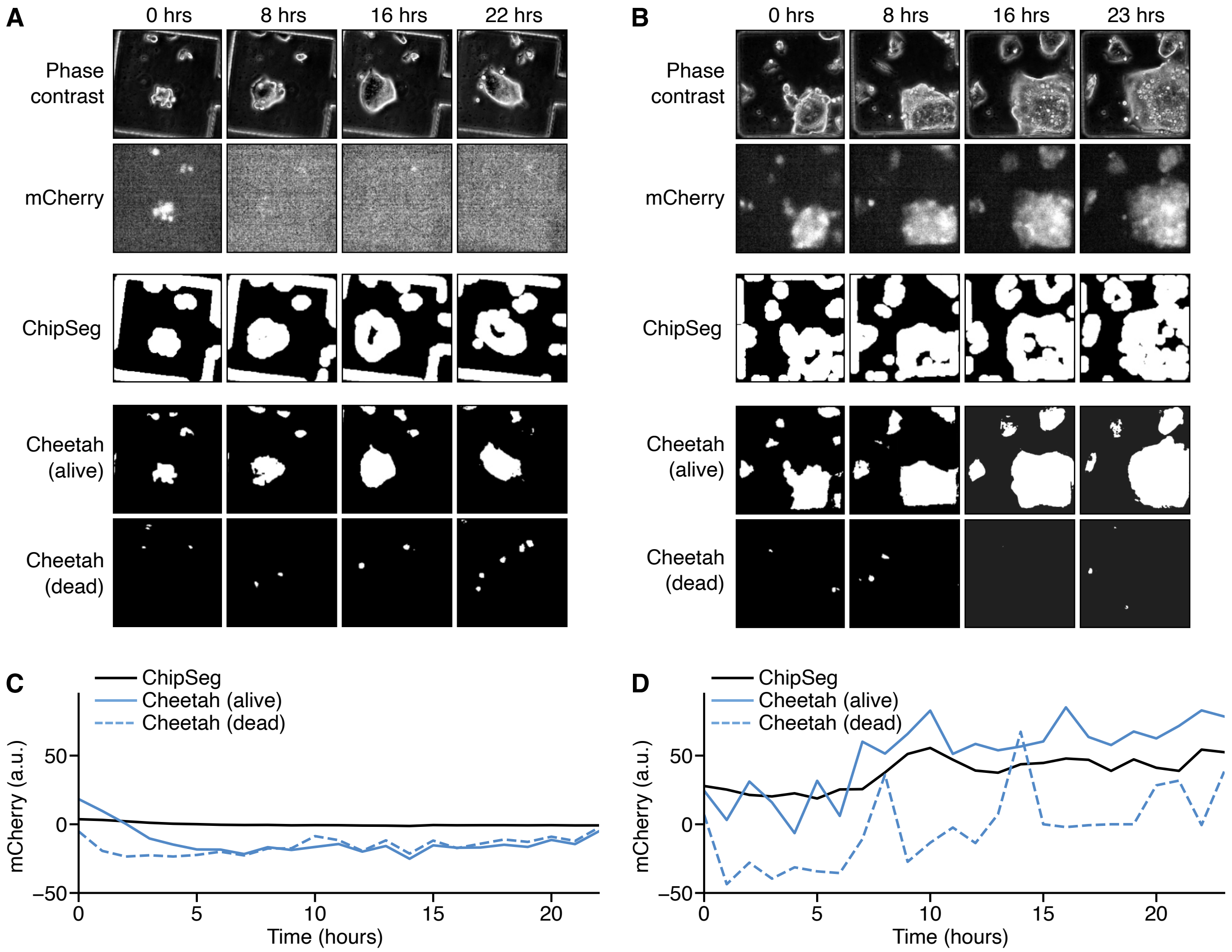


**Supplementary Figure 2: Open-loop protein expression in mammalian cells (replicate 1).** (**A**, **B**) Time-lapse images of mouse embryonic stem cells (mESCs) growing in the system for phase contrast and mCherry fluorescence, as well as segmentation masks for cells generated using ChipSeg and Cheetah in the absence (panel A) and presence (panel B) of Doxy (1 µg/mL) and TMP (100 mM), respectively. (**C**, **D**) Average mCherry fluorescence of the cell segmentation mask over time calculated using either ChipSeg or Cheetah in the absence (panel C) and presence (panel D) of Doxy (1 µg/mL) and TMP (100 mM), respectively.

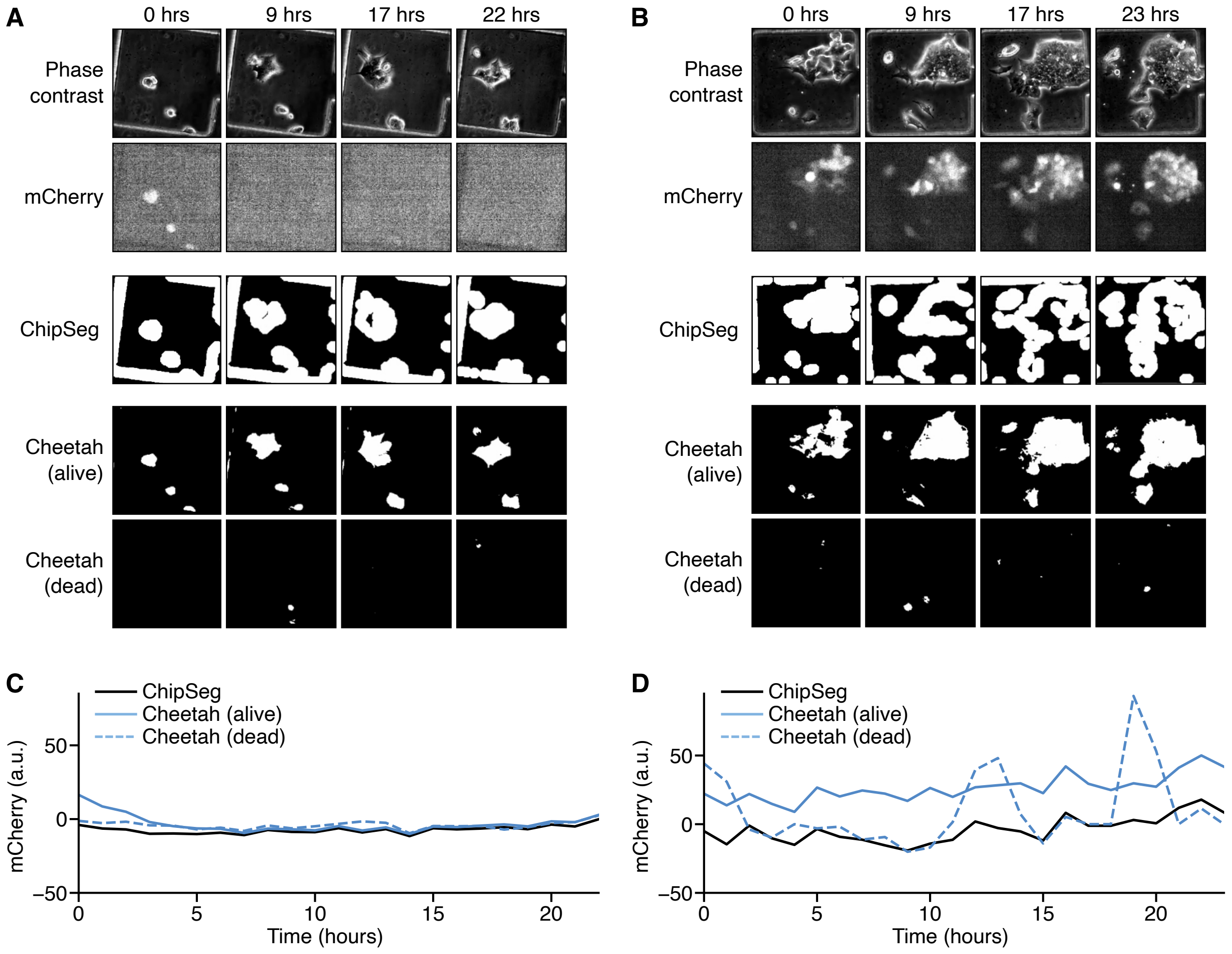


**Supplementary Figure 3: Open-loop protein expression in mammalian cells (replicate 2).** (**A**, **B**) Time-lapse images of mouse embryonic stem cells (mESCs) growing in the system for phase contrast and mCherry fluorescence, as well as segmentation masks for cells generated using ChipSeg and Cheetah in the absence (panel A) and presence (panel B) of Doxy (1 µg/mL) and TMP (100 mM), respectively. (**C**, **D**) Average mCherry fluorescence of the cell segmentation mask over time calculated using either ChipSeg or Cheetah in the absence (panel C) and presence (panel D) of Doxy (1 µg/mL) and TMP (100 mM), respectively.

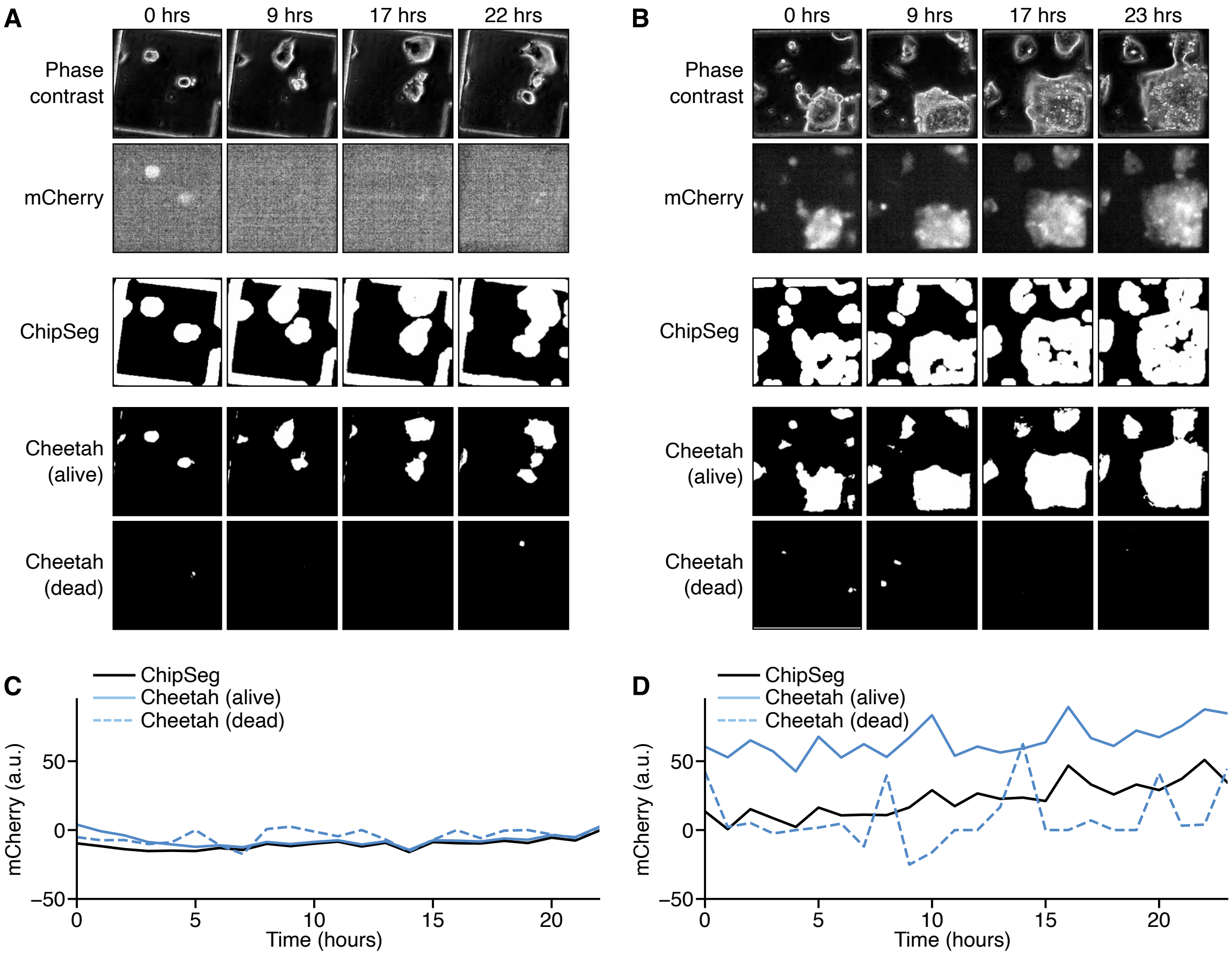


**Supplementary Figure 4: Open-loop protein expression in mammalian cells (replicate 3).** (**A**, **B**) Time-lapse images of mouse embryonic stem cells (mESCs) growing in the system for phase contrast and mCherry fluorescence, as well as segmentation masks for cells generated using ChipSeg and Cheetah in the absence (panel A) and presence (panel B) of Doxy (1 μg/mL) and TMP (100 mM), respectively. (**C**, **D**) Average mCherry fluorescence of the cell segmentation mask over time calculated using either ChipSeg or Cheetah in the absence (panel C) and presence (panel D) of Doxy (1 μg/mL) and TMP (100 mM), respectively.

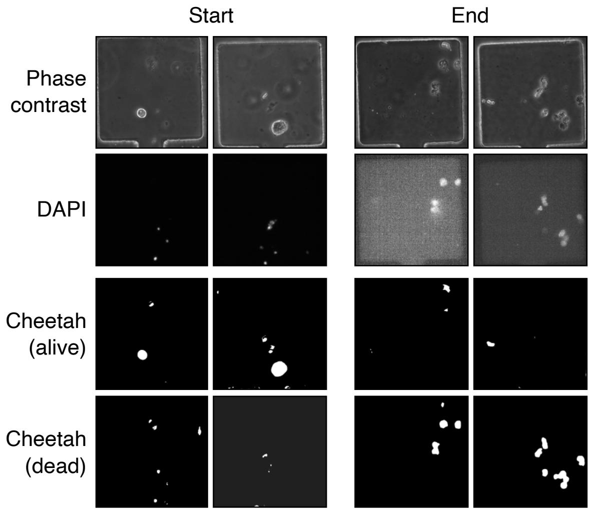


**Supplementary Figure 5: Experimental verification of alive/dead cell classifications.** Images of DAPI stained cells taken at the start and end of a time-lapse experiment. Cheetah segmentation masks for alive and dead cells generated using only widefield images.

**Supplementary Table 1: Integral square errors for time-lapse segmentations**

| Experiment | Cell state | Integral square error (×10⁴) | | | | |
|---|---|---|---|---|---|---|
| | | **Controlled Chamber** | **Uncontrolled Chamber 1** | **Uncontrolled Chamber 2** | **Uncontrolled Chamber 3** | **Uncontrolled Chamber 4** |
| Figure 3 | alive | 1.1 | 5.8 | 4.7 | 1.4 | 16.0 |
| | dead | 1.4 | 4.2 | 8.1 | 1.8 | 26.0 |
| Supplementary Figure 1A,C | alive | 6.1 | 12.0 | 10.4 | 9.1 | – [a] |
| | dead | 8.8 | 15.0 | 11.0 | 14.0 | – [a] |
| Supplementary Figure 1B,D | alive | 5.1 | 12.0 | 7.4 | – [a] | – [a] |
| | dead | 12.0 | 14.0 | 5.1 | – [a] | – [a] |

a. Not all experiments had 4 additional uncontrolled chambers present.

**Supplementary Movie Captions**

**Supplementary Movie 1: Open-loop experiment of bacterial cells.** ChipSeg- and Cheetah-based segmentation results are shown, comparing the computed masks, cell number and GFP fluorescence over time.

**Supplementary Movie 2: Open-loop experiment of mouse embryonic stem cells.** ChipSeg- and Cheetah-based segmentation results are shown, comparing the computed masks and mCherry fluorescence. Cheetah also classify cells as live and dead and provides fluorescent protein dynamics of each.

**Supplementary Movie 3: External feedback control experiment of mouse embryonic stem cells performed using Cheetah-based segmentation.** Offline and online ChipSeg- and Cheetah-based segmentation results are shown, comparing the computed masks cell number and mCherry fluorescence. The control input provided during the experiment and the set-point control reference are also shown.

**Supplementary Movie 4: Detailed analysis dashboard for bacteria growing in a microfluidic chip.** Top two panels on the left show the phase contrast and GFP fluorescence images from the microscope. Top right panel shows detailed analysis of the phase contrast image with cells labelled by colour and a light grey bounding box and their centre of mass and major axis (i.e. orientation) denoted by a red circle and line, respectively. The bottom two panels show the time course of both cell count and single-cell GFP fluorescence (with the average shown as a solid line and ± the standard deviation depicted by the shaded area).
